# Normative modeling of neuroimaging data using generalized additive models of location scale and shape

**DOI:** 10.1101/2021.06.14.448106

**Authors:** Richard Dinga, Charlotte J. Fraza, Johanna M.M. Bayer, Seyed Mostafa Kia, Christian F. Beckmann, Andre F. Marquand

## Abstract

Normative modeling aims to quantify the degree to which an individual’s brain deviates from a reference sample with respect to one or more variables, which can be used as a potential biomarker of a healthy brain and as a tool to study heterogeneity of psychiatric disorders. The application of normative models is hindered by methodological challenges and lacks standards for the usage and evaluation of normative models. In this paper, we present generalized additive models for location scale and shape (GAMLSS) for normative modeling of neuroimaging data, a flexible modeling framework that can model heteroskedasticity, non-linear effects of variables, and hierarchical structure of the data. It can model non-Gaussian distributions, and it allows for an automatic model order selection, thus improving the accuracy of normative models while mitigating problems of overfitting. Furthermore, we describe measures and diagnostic tools suitable for evaluating normative models and step-by-step examples of normative modeling, including fitting several candidate models, selecting the best models, and transferring them to new scan sites.

## Introduction

The goals of normative modeling are to estimate centiles of a distribution of a specific brain phenotype in a population with respect to one or more variables and use these estimates to quantify the biological “abnormality” or deviation of previously unseen individuals from the reference model (Marquand et al. 2016). The resulting abnormality scores can be used as a nonspecific biomarker of healthy brain development or psychiatric disorders at an individual level. This is analogous to the use of growth charts in pediatrics that describe normative ranges of body measurements in children with respect to age and sex. So far, normative models have been shown to be able to detect white matter brain injury in infants (O’Muircheartaigh et al. 2020), discriminate between healthy controls and patients with schizophrenia, bipolar (Wolfers et al. 2018), or ADHD (Wolfers et al. 2020), and have served as a tool to study the heterogeneity of mental disorders (Wolfers et al. 2018); (Marquand et al. 2019). Despite its recent successes, the development of normative models faces many methodological challenges. The ideal method for normative modeling should be able to be used in big datasets, accurately estimate centiles of distributions of various shapes, and handle multiple variables of interest and nuisance variables. The resulting model should be able to be shared with other researchers and should be able to be recalibrated for usage in a new scan site. However, many of the currently used methods meet only some of these criteria, which hinders the applicability of normative models.

Several methods had been recently proposed addressing specific needs of normative modeling.Kia et al. (Kia et al. 2020) proposed a method, which allowed for modeling of heteroscedastic noise, further improving model fit in multi-site datasets. This method also allows for model updating and federated learning. However, since it is based on a posterior sampling inference, it is computationally demanding. Bayer et al. (Bayer et al. 2021) proposed an extension where an effect of age is modeled as a gaussian process regression, which further improved the fit of normative models, however, this method is also computationally very demanding due to its use of a posterior sampling inference and a bad computational scaling of the gaussian process algorithm. Both of these methods also assume a Gaussian distribution of the residuals. Fraza et al. (Fraza et al. 2021) proposed Bayesian regression with likelihood warping, which allows to flexibly fit a non-Gaussian noise distribution. It also does not depend on a posterior sampling for an inference, therefore it is computationally faster and scalable to big datasets. Another way to relax the Gaussian assumption is to use quantile regression (Abbott et al. 2018), where the selected quantiles are modeled directly, without any other distributional assumptions. This method’s drawback is that the derived abnormality scores are discrete and therefore less informative, and the resulting models are often less stable than alternatives, especially for extremely low or high quantiles, where the data are usually the sparsest. There is also a possibility that the estimated quantiles will cross, which would further complicate the analysis. The models are also harder to interpret since every estimated quantile curve corresponds to a separate model.

Here we present an approach to normative modeling using generalized additive models of location, scale and shape (GAMLSS)(Stasinopoulos et al. 2017; Rigby and Stasinopoulos 2005). GAMLSS was used to develop growth charts for the World Health Organization (WHO Multicentre Growth Reference Study Group 2006; Borghi et al. 2006), and it is also suitable for normative modeling of neuroimaging data. GAMLSS is a flexible modeling framework that incorporates the benefits of many previously used methods in one unified package. It allows for the modeling of non-linear effects with respect to many variables, it handles hierarchical models and heteroscedastic models, and it can also fit non-Gaussian distributions. Due to recent algorithmic advancements (Wood and Fasiolo 2017), it is faster than alternatives, including Fraza et al.(2021). The contribution of this paper is threefold; 1) we introduce GAMLSS for normative modeling of the brain. 2) We describe ways to evaluate normative models and their potential pitfalls, which are not limited to GAMLSS. 3) We show step-by-step didactic examples that illustrate how to perform normative modeling, focusing on modeling neuroimaging variables based on age, sex, and scan-site.

## GAMLSS

The GAMLSS framework (Stasinopoulos et al. 2017; Rigby and Stasinopoulos 2005) is an extension of the generalized additive model (GAM) framework (Hastie and Tibshirani 1990; Wood 2017), which itself is an extension of well-known general and generalized linear models (Figure 1). Generalized additive models differ from generalized linear models in a way that the effect of predictors on the response variable is not linear, but it is modeled by a non-linear smooth and a possibly non-parametric function. GAMLSS are like GAM models, but they can also be used to model multi-parameter distributions, where each parameter of the distribution, i.e., location, scale, and shape, can be modeled as a function of several input variables, thereby accommodating a wide range of distributions. For example, most linear models assume homoskedasticity and Gaussianity of the residuals. We can use GAMLSS to model both location and the scale of the Gaussian distribution based on several predictors, thus relaxing the homoskedasticity assumption. We can also use GAMLSS to model a four-parameter distribution such as Sinh-Arcsinh (SHASH) (Jones and Pewsey 2009) instead of Gaussian distribution, where the parameters of the distribution corresponding to location, scale, skewness, and kurtosis of the distribution can all be modeled as a function of several input variables. This allows us to accurately estimate centiles of the distribution even when the residuals are not normally distributed.

**Figure 1.**
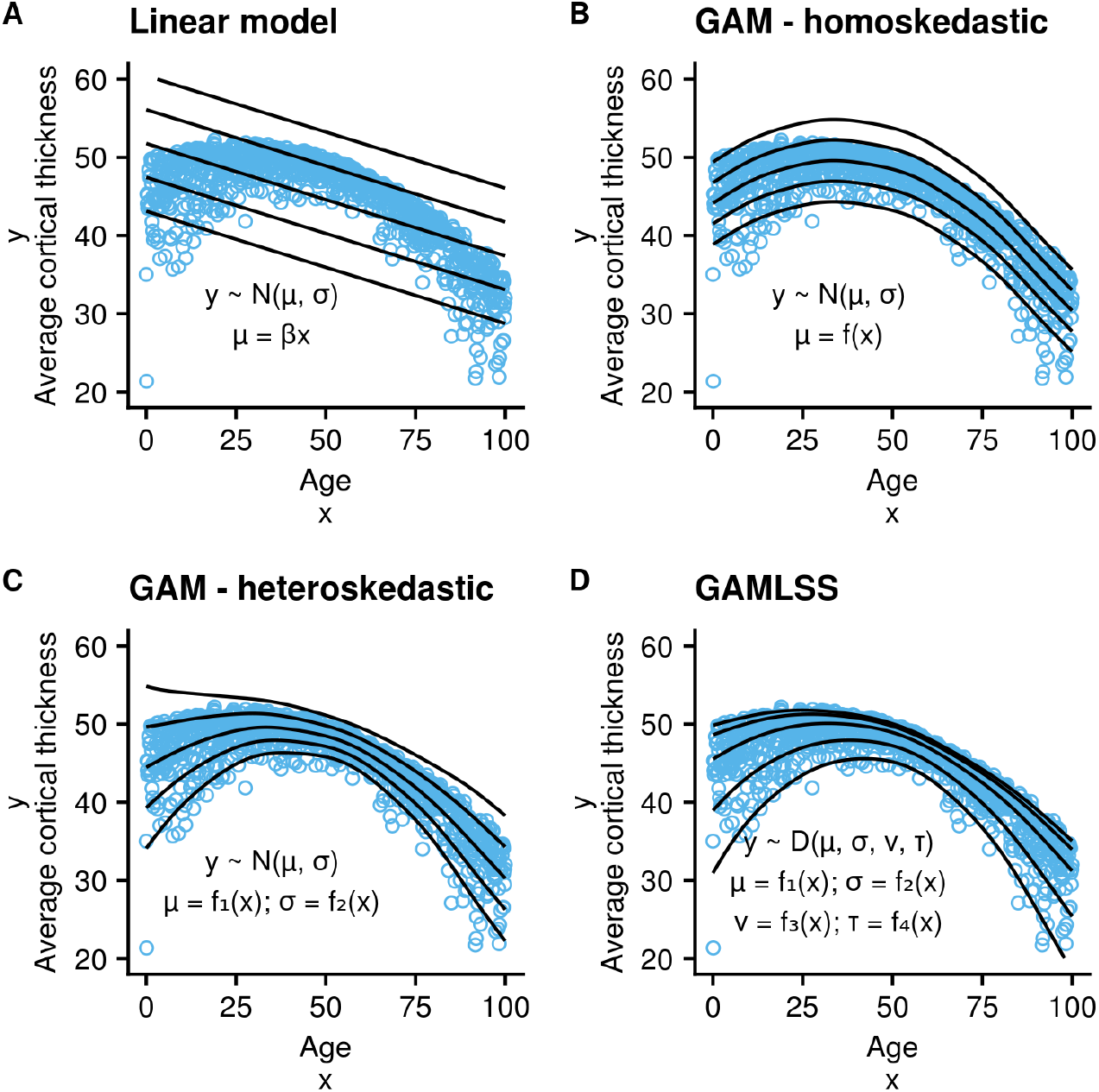
From a linear model to GAMLSS: **A:** Linear model models mean the target variable as a linear function of x. **B:** generalized additive model (GAM) models the mean of the target variable as a smooth function of x. **C:** GAM - heteroskedastic models mean and variance of the response variable as functions of x. **D:** GAMLSS allows to model y as a multiparameter non-gaussian distribution, where location, scale, and shape parameters can be modeled as functions of input variables.

### From a linear model to GAMLSS

A linear model can be described as

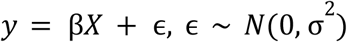

or using a slightly different notation

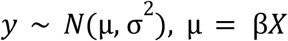

where y is the target variable, βis a vector of estimated coefficients, X is a design matrix and ϵ is an error vector, i.e., y depends linearly on the predictors X and the error term is assumed to be normally distributed. GAM extends LM by relaxing the linearity assumption:

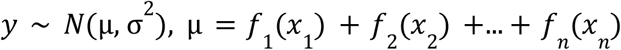

Where *f*(*x_i_*) is a possibly non-linear or non-parametric function of *x_i_*. The original formulation of GAMs (Hastie and Tibshirani 1990) allowed for almost arbitrary functions, including loess smoothing, decision trees, or neural networks that could then be an additive part of a larger model. More modern interpretations of GAMs (Ref: Wood) limit these functions to smooth mostly spline-based functions of X, where the complexity of the model can be controlled by penalizing the wiggliness of a function (In GAM nomenclature, ‘wiggliness’ is given a precise technical meaning usually referring to an integrated second derivative of a function). The wiggliness of the function is a parameter that can be estimated from the data, which allows for flexible modeling of non-linear effects with an automatic selection of model complexity. In a linear model, we can model non-linear effects by a spline basis expansion of the input variables, and the number of knots will control the model complexity. In GAM, the number of knots is usually fixed at some high number, and the model complexity is controlled by directly penalizing the second derivative (i.e., the wiggliness) of the function, which can be implemented as a quadratic penalty on weights of the spline basis expansion.

GAMLSS is an extension of GAM introduced by (Stasinopoulos et al. 2017; Rigby and Stasinopoulos 2005), which allows to model multiple parameters of a multiple parameter distributions as additive functions, i.e., instead of just modeling the expected mean of the target variable and assuming that the distribution around this mean is normally distributed, we can model location, scale, and shape parameters of the distribution. So for a four-parameter distribution

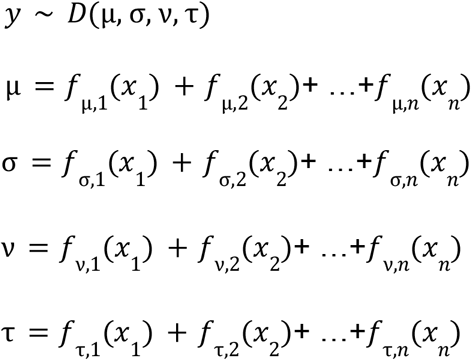

Here D is a parametric distribution with parameters μ, σ, ν, τ, and each of the parameters can be modeled as a separate function of input variables. This allows for accounting for heteroskedasticity by directly modeling the scale of a distribution (σ) as a function of other variables or modeling shapes of non-gaussian distributions. Each parameter can be modeled based on a different set of variables and using a completely different structural form. So it is possible to have a complex model for μ but at the same time a very restricted model for σ.

GAMLSS is usually estimated using a penalized maximum likelihood estimation. The smoothing functions *f*(*x*) can be expressed as *Z*_γ_ where*Z*is a matrix of basis expansion of *x*, and γ is a set of coefficients penalized using a quadratic penaltyλ_γ_^*T*^*G*_γ_ where λ is a smoothing hyperparameter. We can also say thatγare random effects, assumed to be distributed as γ ~ *N* (0, *G* ^−^). If we are estimating a model for 4 parameter distribution, that depends on J smooth functions, the criterion for the penalized maximum likelihood is

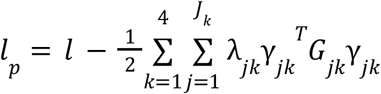

where

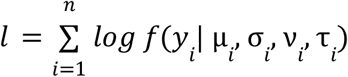

Traditionally, this is done iteratively using a Newton-Raphson or Fisher scoring (Rigby and Stasinopoulos 2005), with recent algorithmic improvements improving computational complexity of these models and allowing their use in big datasets (Wood and Fasiolo 2017). Full Bayesian inference using a posterior sampling is also possible (Umlauf, Klein, and Zeileis 2018).

In the rest of this paper, we are using SHASH (Jones and Pewsey 2009) distribution as implemented in the mgcv library (Wood 2017) as an example multi-parameter distribution that can be fitted using the GAMLSS approach. The four parameters roughly correspond to location, scale, skewness, and kurtosis of the distribution. SHASH has the normal distribution as its special cases, but based on its parameters, it can also be skewed or more or less kurtic than the normal distribution.

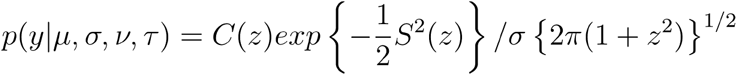

where *C*(*z*) = 1 + *S*(*z*)^2^, *S*(*z*) = *sinh* {*τsinh*^−1^(*z*) – *ν*},and *z* = (*y* – *μ*)/(*στ*) Parameters μ, σ, ν, τcontrol location, scale, skewness, and kurtosis respectively.

## Model diagnostics

For the normative modeling approach, it is important to check if the data are reasonably well approximated by the model, even more so than for a more traditional hypothesis-testing approach. Since the normative modeling goal is to estimate an individual’s abnormality scores based on the mean and centiles of the distributions, the deviations from model assumptions are much more severe as they will bias the estimated subject’s abnormality scores. The GAMLSS framework is flexible enough to relax many assumptions, mainly linearity, homoskedasticity, and Gaussianity of residuals. However, to appropriately adjust a model, it is important to detect these problems and monitor if different modeling choices improved model performance. This section describes selected measures and graphical tools for model diagnosis and validation of normative models.

All these measures can be calculated in the training set or the test set. We can expect a more complex model to always perform seemingly better in the training set than in a test set. However, we recommend to perform model diagnostics in both the training set and test set since even with this bias, many potential problems of model fit could be reliably detected using the training data. Although the model improvement should be interpreted with caution, and validation on the training set should not be a replacement for validation in the test set.

### Goodness of fit

The most important measures are the measures of total goodness of fit that are sensitive to all possible misspecification of the model, including badly estimated scale or shape of the distribution, heteroskedasticity, or differences between linear or nonlinear models. For this reason, various popular measures used to validate regression models, such as mean absolute error (MAE), R^2^, correlation, or root mean squared error (RMSE), are not appropriate since these are calculated based on the predictive meanbut do not take into account other aspects of modeled distribution. Appropriate measures are so-called proper scoring rules (Gneiting, Balabdaoui, and Raftery 2005), which are maximized by the true data-generating model. One possible measure is the logarithmic score or validation global deviance (Stasinopoulos et al. 2017) or sometimes predictive deviance.

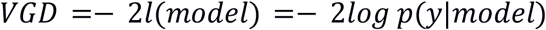

where *l*is the model likelihood, and *p*(*y*|*model*) is a predicted probability density by the model, at the target variable y in the test set. It is called a “validation global deviance”, since the model is estimated in a training set, and the deviance is calculated based on the data in the test set. The logarithmic score is defined similarly

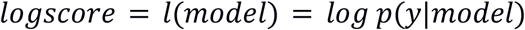

which can be scaled to aid interpretation as for example, deviance explained or mean standardized log loss (MSLL) (Rasmussen and Williams 2005), which is averaged across all data points. The MSLL is computed as follows

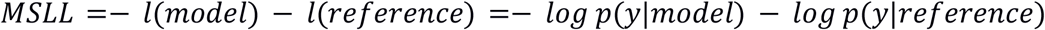

Where *reference* is a reference model, usually a trivial model that predicts only an mean and standard deviation of the sample. We can see that these measures are just linear transformations of each other, therefore the relative ranking of models will be the same, provided the reference model used to calculate MSLL will not change. The deviance is the predicted density of an individual’s observed value, and therefore these measures depend on the location, scale, and shape of the predicted distribution.

One potential drawback is that these measures are sensitive to outliers, and unless care is taken with regard to treatment of outliers, this might lead to selecting the wrong model. For example, removal or Winsorizing of outliers will lead to models with underestimated variance and kurtosis of the distribution. One way to make these measures more resistant to outliers while keeping them proper is to use a censored likelihood (Diks, Panchenko, and van Dijk 2011), which modifies likelihood definition to

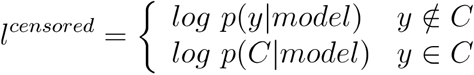

where

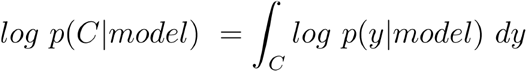

where C is a chosen censored region. So if a predicted density is not in the censored region, the likelihood is calculated, as usual, otherwise, the log probability of a y being anywhere in the censored region given the model is used instead. This censored likelihood can be used instead of standard likelihood in other likelihood based measures (e.g., VGD or MSLL mentioned above), to create their robust versions.

For example, we might decide to censor values that are not between the 1st and 99th percentile. If the target value y for a subject is anywhere between the 1st and 99th percentile, the usual logarithmic score will be calculated. If the y is anywhere else, regardless of if the subject is an extreme outlier or not, the resulting score will be log(0.02), since the probability of y being outside of the 1st-99th percentile range is 2 percent. This means that this score would prefer models where a shape of the distribution between the 1st-99th percentiles is modeled accurately, and 2% of subjects are outside of this interval, but the shape of the distribution outside of the interval would not matter. The censored likelihood can also be used if we want to evaluate if some specific part of the distribution is modeled correctly, for example, only the distribution’s right tail. For example, if we want to detect a disorder, and we know that the disorder has a neurodegenerative effect, we might want to focus on accurate modeling of the right tail of the distribution since this is where we expect clinical cases to lie. Although likelihood based measures are affected by all aspects of model fitting, it is still worth validating specific aspects of model fitting such as calibration and central tendency separately.

### Central tendency

Measures of goodness of fit of the central tendency measure how well the center of the distribution is predicted. This is mainly a concern when comparing which model is better at modeling a possibly non-linear trend in the data. If the central tendency is not fitted correctly, we might expect, for example, that z-scores for younger and older subjects are systematically underestimated, and middle-aged subjects are overestimated. The center of the distribution can refer to the mean, the median, or the mode of the predicted distribution, and different interpretations may be preferred, for example, depending on whether the distribution is symmetric. The measures include the traditional measures commonly used to evaluate the performance of regression models, such as MAE, correlation, R2, or RMSE. There is nothing inherently better or worse about these measures; they all have their place, however, there are a few potential pitfalls when evaluating non-Gaussian models. In a Gaussian model, mean, median, and mode are the same, and they are fitted to minimize MSE. This means that MSE will often prefer Gaussian models, even if non-gaussian models perform better. The safest option is to use the predicted median and evaluate it using MAE, which is minimized by the median (Hanley et al. 2001). The predicted mean might be technically hard to compute since if the mean is not one of the estimated parameters of a distribution, we would have to numerically integrate the whole density function to calculate it. Measures based on mean might also be negatively affected by skewness or outliers in the data.

### Calibration

Calibration assesses if the estimated centiles of a distribution match the observed centiles, or if the modeled distribution’s shape is well approximating the observed distribution. Calibration in normative models is best evaluated using quantile randomized residuals, also known as randomized z-scores or pseudo-z-scores (Umlauf, Klein, and Zeileis 2018; Dunn and Smyth 1996). These are quantiles of the fitted distribution mapped to z-scores of a standard normal distribution corresponding to the same quantiles. So if a subject is in a 95th percentile according to the GAMLSS model, their randomized z-score will be 1.97 (because 95 percentile of a standard normal distribution is at 1.97), regardless of the shape of the modeled distribution.

If the modeled distribution matches the observed distribution well, the randomized z-scores should be by construction normally distributed, regardless of the modeled distribution’s shape. The most informative are visual displays of the residuals in the form of either histogram, Q-Q plots, P-P plots, or wormplots (detrended Q-Q plot) (Hanley et al. 2001; van Buuren and Fredriks 2001). Various ways to visualize deviations from the assumed distribution are shown in figure 2. Of course, visual inspection is not always possible, for example, when modeling hundreds or even hundreds of thousands of voxels, therefore we need to have quantitative summary measures as well. Since the pseudo-z-scores should be normally distributed, calibration can be measured as various forms of deviations from the normality of the distribution pseudo-z-scores. One possibility is to use the test statistics from the Shapiro-Wilk test of normality (Shapiro and Wilk 1965) or a test statistic from other normality tests. In this case, we are not interested in calculating a p-value from the test but in using the test statistics to quantify the degree to which the observed distribution deviates from the assumed normal distribution. W is usually close to 1. Perfectly normally distributed data would obtain the value of W of 1, while the values less than 1 indicate deviation from normality. We can also measure scale, skewness, and kurtosis of the pseudo-z-scores separately to give us more insight into how the model is potentially misspecified.

**Figure 2:**
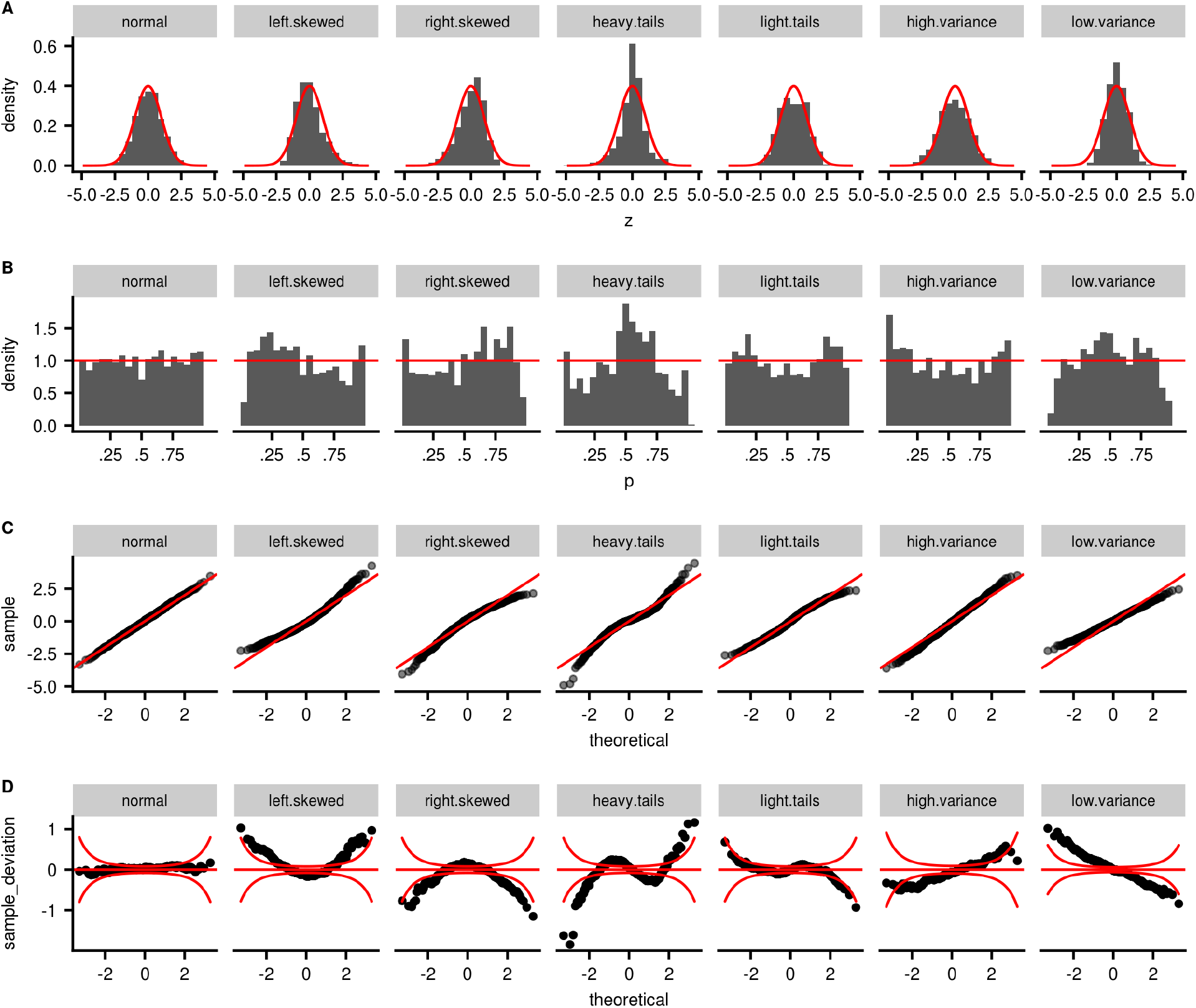
different ways how a model can be miscalibrated and how to visually assess it using multiple methods. **A:** histogram of pseudo-z-scores, which if a model is well-calibrated should follow a normal distribution with mean 0 and standard deviation 1. **B:** Histogram of estimated percentiles, which should be uniform for well-calibrated models. **C:** Q-Q plot comparing theoretical quantiles of a normal distribution with mean 0 and standard deviation 1 to observed quantiles. **D:** worm plot (van Buuren and Fredriks 2001), which is constructed from Q-Q plot, but instead of observed quantiles on the y-axis, it has the difference between observed and theoretical quantiles. The red confidence band shows a region where approximately 95% of points should be if a model is well calibrated. In all cases, the model assumes that the distribution is normal with a mean 0 and standard deviation of 1. The first column shows an example of when this assumption is correct. Columns 2 and 3 show different visualizations of a skewed distribution, columns 4 and 5 show leptokurtic and platykurtic distributions, and columns 6 and 7 show normal distributions but with a larger and smaller estimated variance relative to the true variance.

### Heteroskedasticity

Heteroskedasticity means that a variance of the response variable changes as a function of some variable in the dataset, for example, with age or scan-site. If this aspect is not modeled correctly, some subjects’ abnormality will be systematically overestimated, and others underestimated. To investigate heteroskedasticity, we can again examine randomized residuals, which should by construction have a standard deviation of 1, the standard deviation should not be related to any variables in the data, i.e., in each scan-site or age bin. There are many ways how heteroskedasticity can be detected. One possibility is to examine the data visually by plotting squared residuals versus predicted variables and fitting a curve to this plot. If the residuals are homoskedastic, this curve should be close to a straight line. We can also calculate the goodness of fit of this curve to use as a quantitative measure of heteroskedasticity.

## Model comparison and model selection

The highest standard for model validation is to validate the models in the independent test set since, in the training set, more complex models would be likely preferred due to over-fitting. However, there are also several ways to compare models in the training set. First, if the models are nested, we can use a likelihood ratio test, or if they are not nested, we can use AIC or BIC, or more general GAIC (ref). In our experience, these methods work well for simple models, but they do not reliably select the best model in GAMLSS models with many smooth effects, random effects, and multiple parameters of a distribution to be fitted. Therefore, we recommend using them as imperfect heuristics and not as a replacement to a more thoughtful model selection process.

Importantly, the AIC and BIC values are not by themselves interpretable and should be interpreted relative to AIC and BIC values from other candidate models (Burnham and Anderson 2010)(Wagenmakers and Farrell 2004). To guide this interpretation, we can compute for the ith model

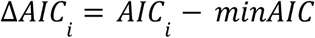

where minAIC is the smallest AIC from all candidate models. From ΔAIC we can calculate the relative AIC

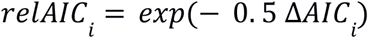

Which is interpreted as a relative likelihood of the best model and the ith model. From the relative AIC we can calculate Akaike weights which are obtained by normalizing relative AIC by the sum of relative AIC of all candidate models

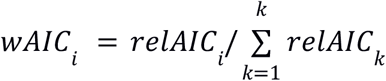

Akaike weight is interpreted as the probability that a model is the best model from a set of candidate models (Burnham and Anderson 2010)(Wagenmakers and Farrell 2004). The same measures can be calculated for BIC as well, but keep in mind these are just heuristics that might not perform well in every situation.

Cross-validation is the safer option when it is important to estimate which model will perform better out of the sample accurately. One useful visualization for model comparison is to plot estimated randomized z-scores from one model against estimated randomized z-scores from another model. Often the z-scores are not practically different even though the quantitative measures might suggest they are. In this case, it is not important to select the best model since the differences in predictions might be minimal with a little effect on subsequent analysis.

## Transfer to a new site

Sometimes we might want to use a model in a new site where the data from this site were not available to us during the training process. In this case, we will not know what the site effects should be, and therefore the estimated z-scores might be biased, often severely. It is possible to use a small subsample of subjects from the new site to recalibrate parts of the model to better fit the new site’s distribution while keeping the rest of the model fixed.

A simple way would be to re-estimate the intercept of the new site while keeping the smooth effect of age fixed. In a gaussian, homoskedastic model, this can be performed by

1. select a small subset of subjects from the new site for recalibration
2. calculate residuals for the selected subjects, according to the original model, i.e., subtract the predicted values from the observed values
3. estimate the intercept corresponding to the new site as the intercept of the original model plus the mean of the residuals
4. the standard deviation (sigma) corresponding to the new site can be calculated as a standard deviation of these residuals

This simple procedure would change the intercept and standard deviation for the model to correspond better to the data in a new site while keeping the estimated non-linear effect of age the same as in the original model. We might also additionally improve the fit by putting a prior over these estimates. However, this method might not work well enough for heteroskedastic or non-gaussian models, because 1) predicted values from the model and an intercept of the model usually do not correspond to an average of the predicted distribution, so adjusting the intercept by an average of residuals will not produce correct results. 2) the sigma parameter in a SHASH distribution (Jones and Pewsey 2009) is not the standard deviation; the standard deviation depends on other parameters of the distribution as well, so adjusting sigma by the standard deviation of the residuals would also not produce correct adjustment.

Instead of making adjustments based on residuals, we can refit the selected parameters of a model directly while keeping others constant. This can be performed by fitting the new model with predicted parameters of the distributions as input variables, with their coefficients fixed at 1 So, for example, if the model we want to transfer to a new site is

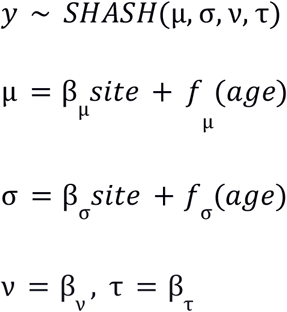

where the site is a matrix of site indicators. In this example, we want to keep all the parameters of the model the same, except β_μ_ and β_σ_. We can fit a new model as follows

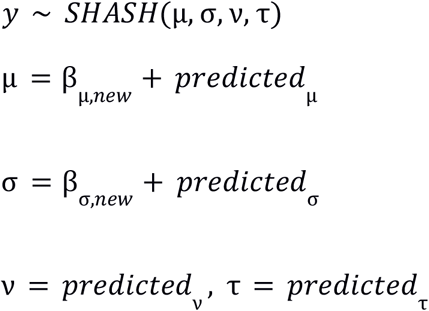

Where onlyβ_μ,*new*_ and β_σ,*new*_ are estimated, and predicted_mu, predicted_sigma, *predicted*_ν_, and *predicted*_τ_ are predicted parameters of the SHASH distributions, and coefficients of these predictors are not estimated in this model but are fixed to 1. In R, this can be performed using an ‘offset’ function. See figure 3.

**Figure 3:**
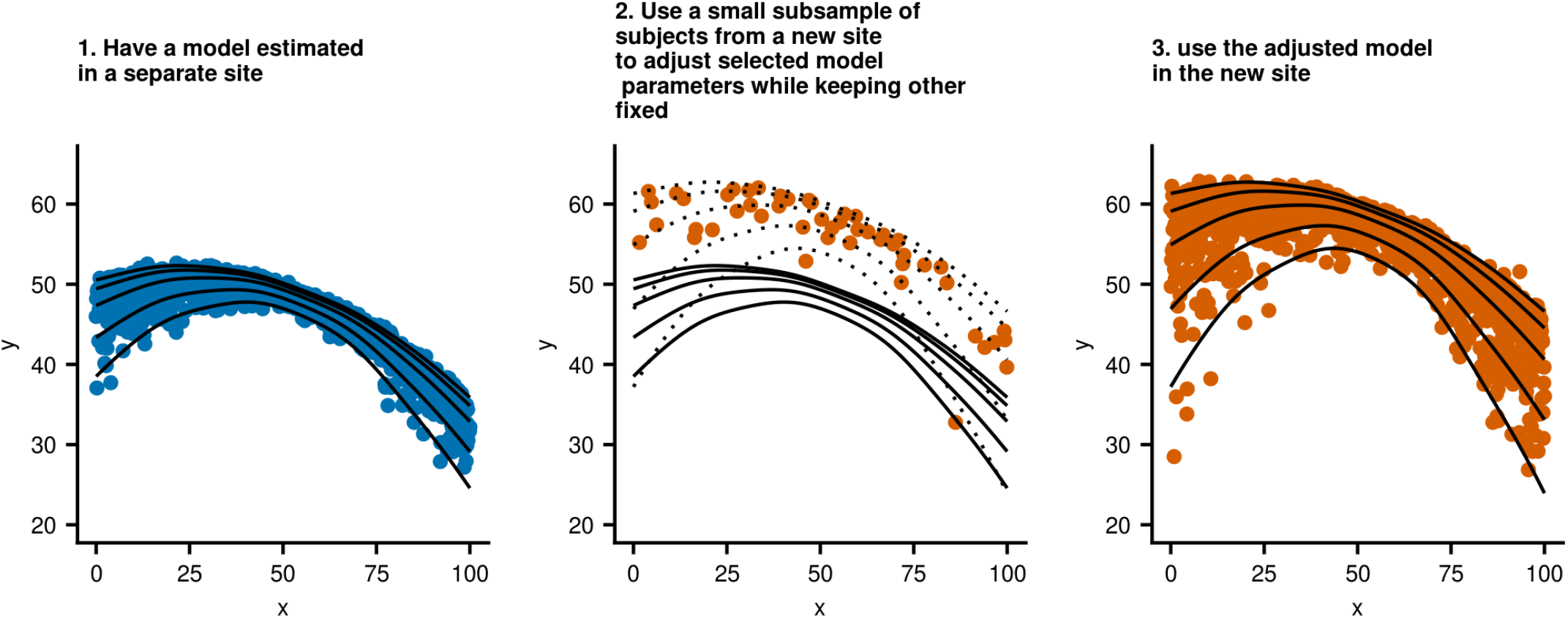
a schematic illustration of a model adjustment to a new site that was not available during the initial model training process. Here, we adjust an intercept and a scale of the model while keeping the estimated non-linear effect, heteroscedastic effect, and non-gaussian shape unchanged.

## GAMLSS example analysis

In this section, we will show a didactic and step-by-step usage of GAMLSS in various neuroimaging settings. We will mainly focus on modeling age and site effects, although the lessons learned should be transferable to other situations.

### 1. modeling the age effect

In the following section, we will use data from UK Biobank UKB (Miller et al. 2016). We used regional cortical thickness data obtained using a processing pipeline as described elsewhere (Alfaro-Almagro et al. 2018). For simplicity, we will start by modeling the effect of age on median cortical thickness in women. We split data into 70% training set and 30% test set. If this was a smaller dataset, we might consider using cross-validation instead of one test set. We start by fitting several models: a Gaussian model with linear effects (M1a), a non-linear Gaussian homoskedastic model (M1b), a non-linear Gaussian heteroskedastic model (M1c), and a non-linear heteroskedastic SHASH model (non-Gaussian)(M1d), and a non-linear heteroskedastic SHASH model where location, scale, and shape also depends on age (Mm1e),:

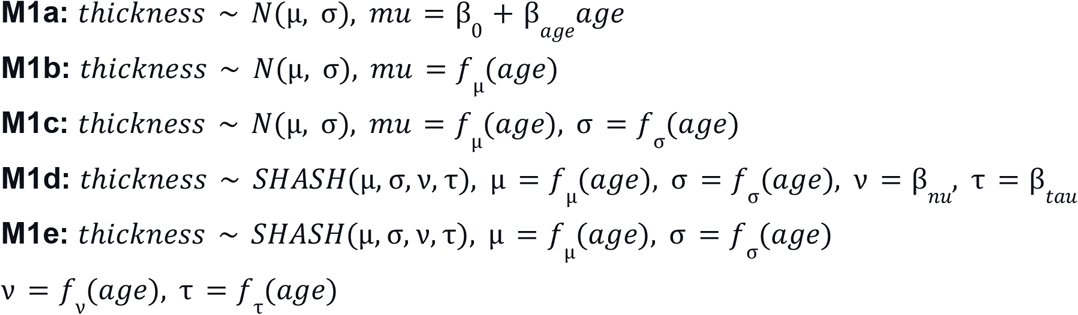

It is recommended to consider models with increased complexity in the order specified above, and not, for example, to consider models that have more complicated models for shape parameters than they have for scale or location (Stasinopoulos et al. 2017). Therefore, we are not going to fit a model with for example a linear effect of age on μ but non-linear on σ. Figure 4 shows resulting model fits, and table 1 contains test set deviance and training set AIC and BIC. We can see that the effect of age seems to be slightly non-linear, while the distribution is slightly heteroskedastic and non-gaussian. This is confirmed by model fit and diagnostics measures. Models M1d and M1e seem to perform the best. Since the differences between the predictions from these models are minimal, we will choose the simpler model, i.e. M1d.

**Figure 4:**
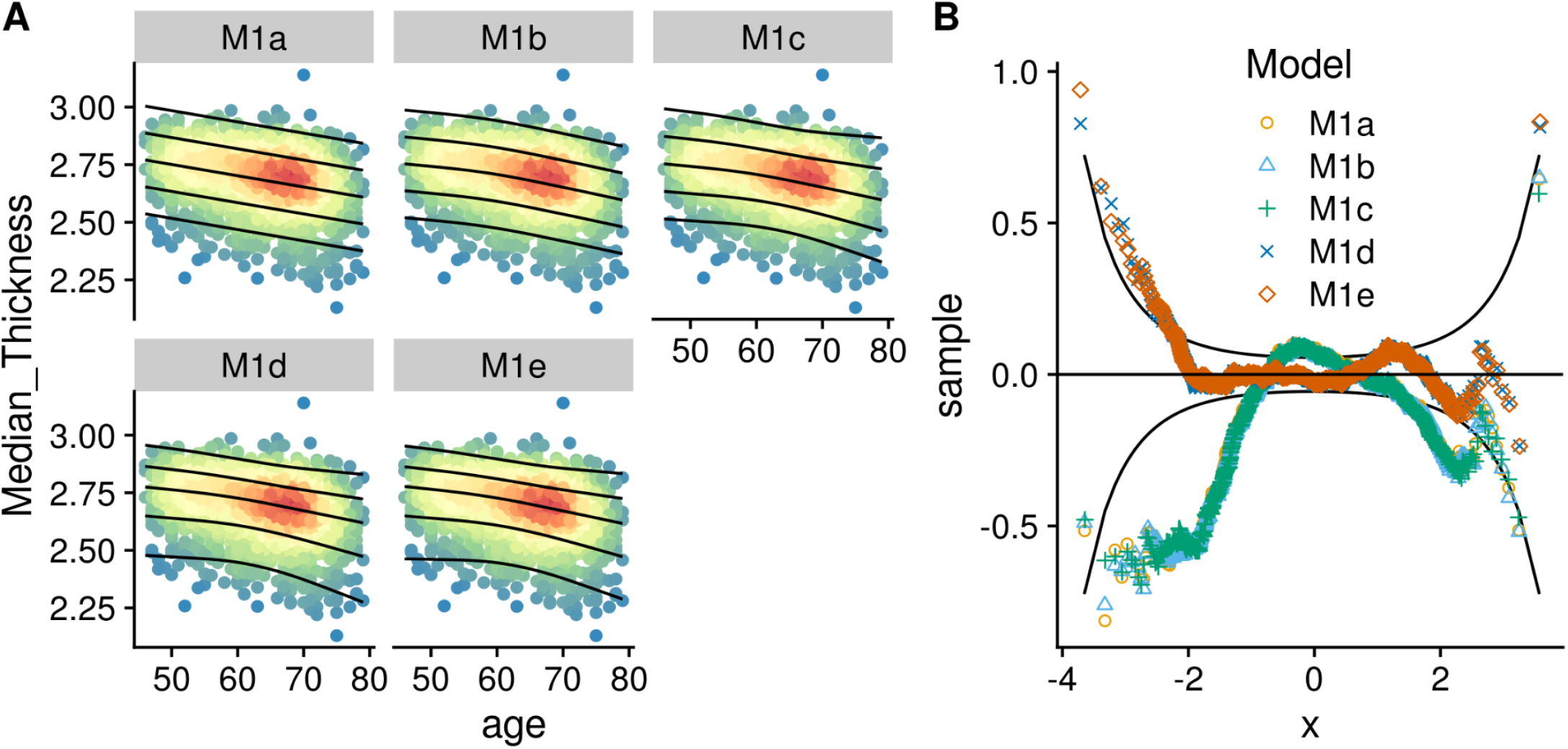
visualization of **A:** estimated quantiles and **B:** model fit of the distribution using a wormplot (detrended Q-Q) plot. The closer the points are to the horizontal line, the better calibrated the model is. The confidence band shows where 95% of the points from a perfectly calibrated model are expected to lie. We see a small non-linear and heteroskedastic effect in the data. According to the wormplot in panel B, models m1d and m1e have a better fit than alternative models but are very similar, seen as mostly overlapping points.

**Table 1a:** AIC and BIC derived measures for different models of age.

| model | AIC | $\Delta AIC$ | AIC weights | BIC | $\Delta BIC$ | BIC weights |
| --- | --- | --- | --- | --- | --- | --- |
| M1a | -5422.29 | 0 | 0 | -5403.23 | 0 | 0 |
| M1b | -5429.8 | 0 | 0 | -5399.78 | 0 | 0 |
| M1c | -5444.86 | 0 | 0 | -5397.28 | 0 | 0 |
| M1d | -5857.19 | 0.38 | 0.38 | -5801.13 | 1 | 1 |
| M1e | -5858.21 | 0.62 | 0.62 | -5788.83 | 0 | 0 |

**Table 1b:** quality of model fits in the test set.

| model | logscore | skewness | kurtosis | W |
| --- | --- | --- | --- | --- |
| M1a | 0.69 | -0.55 | 0.52 | 0.98 |
| M1b | 0.69 | -0.54 | 0.51 | 0.98 |
| M1c | 0.7 | -0.57 | 0.53 | 0.98 |
| M1d | 0.73 | 0.14 | 0.01 | 1 |
| M1e | 0.72 | 0.15 | -0.01 | 1 |

Performance of various models measured in the training set (1a) and the test set (1b); According to a log. Score and normality of pseudo-z-scores. Avg. log. Score: bigger is better, skewness: closer to 0 is better (no skewness), kurtosis: closer to 0 is better, higher than 0 implies heavy tails. W: is a test statistic from the Shapiro-Wilk normality test. Closer to 1 means data pseudo-z-scores are more normal. Both AIC and BIC chose the best model.

### 2. modeling the sex effect

Here, we will examine different ways how the effect of sex on the cortical thickness can be modeled. We can choose not model sex at all (M2a, same model as M1d, but fitted in a sample containing both men and women), choose to model separate intercepts for men and women, while keeping the trajectories the same (m2b), choose to model different age trajectories for men and women (m2c), and last, choose to model all aspects of the model separately for men and women, which is the same as having two separate models (m2d):

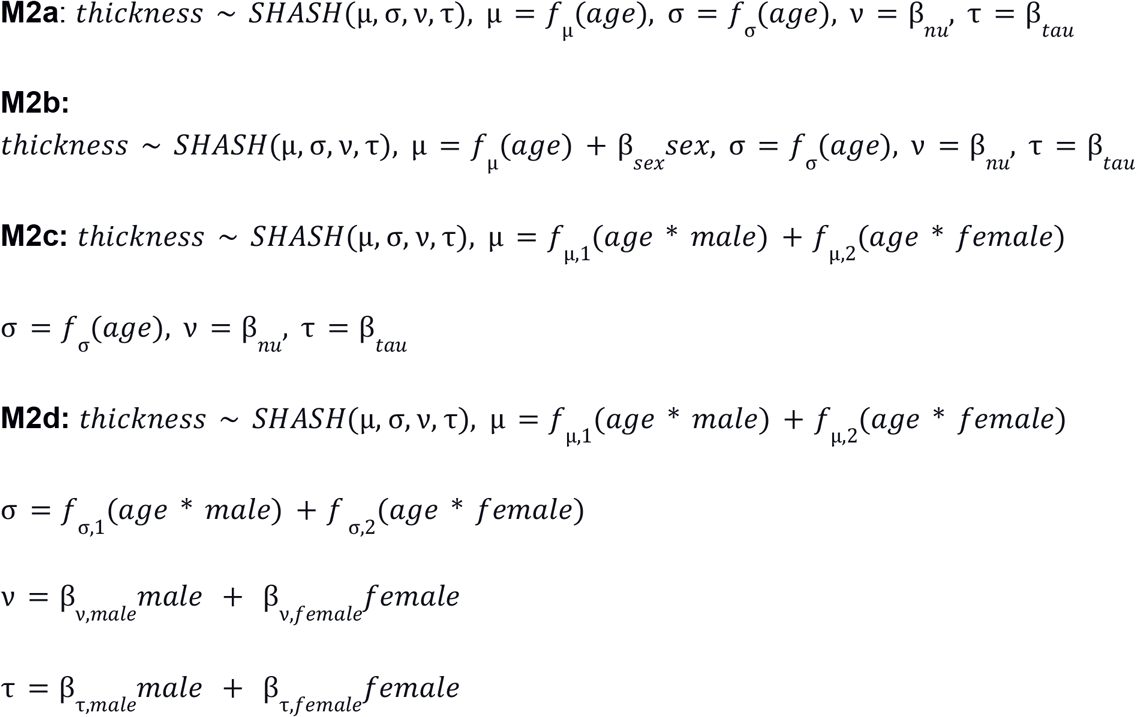

We could also make more complicated models by, for example, modeling the effects of age on ν and τ parameters, but we showed in the previous section that this is not beneficial in this case. Results can be seen in table 2. The results show that sex effects are minimal. In the next section, we are therefore not going to model specific sex effects.

**Table 2a:** AIC and BIC derived measures for different models of sex.

| model | AIC | $\Delta AIC$ | AIC.weights | BIC | $\Delta BIC$ | BIC.weights |
| --- | --- | --- | --- | --- | --- | --- |
| M2a | -12601.6 | 0 | 0 | -12532.4 | 0.23 | 0.23 |
| M2b | -12611.03 | 0.03 | 0.03 | -12534.84 | 0.77 | 0.77 |
| M2c | -12615.79 | 0.29 | 0.29 | -12525.72 | 0.01 | 0.01 |
| M2d | -12616.57 | 0.43 | 0.43 | -12519.73 | 0 | 0 |
| M2e | -12615.5 | 0.25 | 0.25 | -12509.33 | 0 | 0 |

**Table 2b:** quality of a model fit in the test set.

| model | logscore | skewness | kurtosis | W |
| --- | --- | --- | --- | --- |
| M2a | 0.75 | 0.13 | 0 | 1 |
| M2b | 0.75 | 0.13 | 0 | 1 |
| M2c | 0.75 | 0.13 | 0 | 1 |
| M2d | 0.75 | 0.13 | 0 | 1 |
| M2e | 0.75 | 0.13 | 0 | 1 |

All models performed similarly out of sample, indicating a very small sex effect in this data modality.

### 3. modeling site effects

To showcase modeling of site-effects, we use a combined multisite dataset as in (Kia et al. 2020). This dataset includes data from various publicly available studies. The processing details can be found in Kia et al. (2020). As in previous examples, we will model median cortical thickness data obtained using freesurfer 5.3 and 6.0. For simplicity, and due to results from previous section, we do not explicitly model sex effects, although combining modeling of site-effects and sex effects is also possible.

We compare different ways of modeling effects of scan-site. We can choose not to do anything (M3a), model site intercepts for μ as a fixed effect (m3b) or a random effect (m3c), and model site effect for μ and σ as fixed (m3d) or random effects (m3e). We could also model site-specific smooth terms, which could be potentially pooled towards a common trend as in random effect modeling of intercepts see (Pedersen et al. 2019).

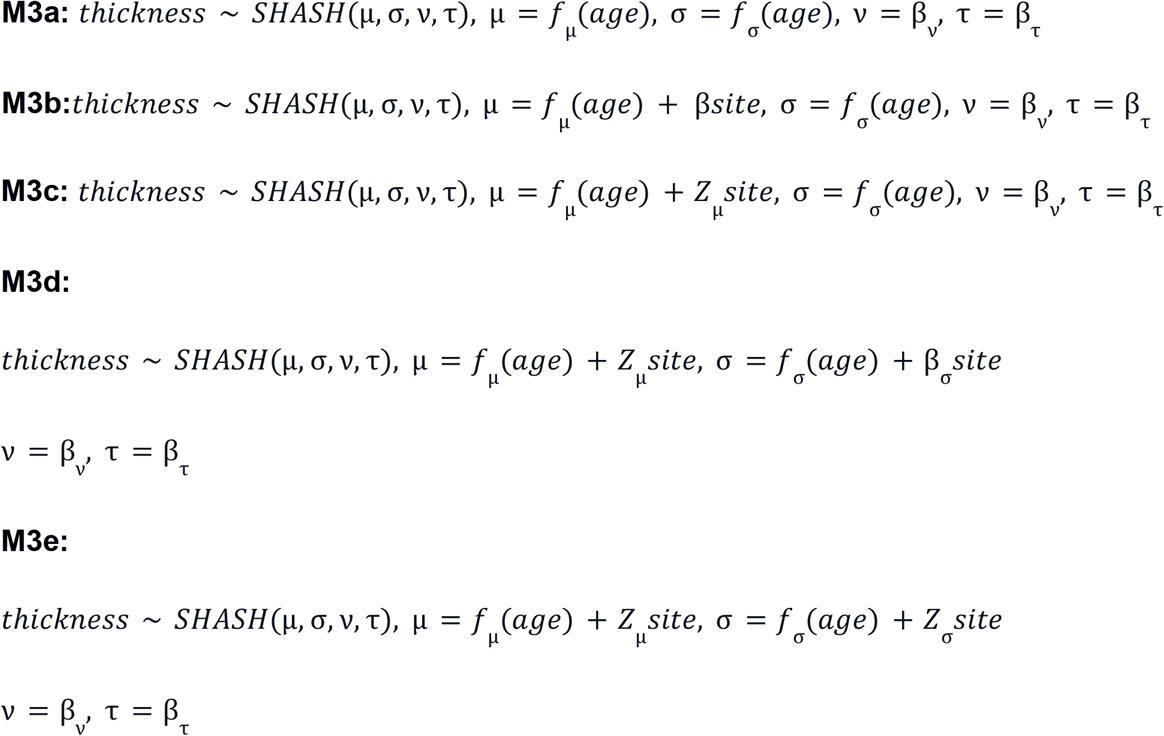

Results can be seen in table 3. We can see that modeling site effects for μ and σ both significantly improve predictions in the new subjects. This can be seen in the deviance, but also when looking at z-scores stratified by site. We can also see that if the site effects are modeled, there is a smaller site effect present in the pseudo-z-scores in the test set. There will always be some site effect noticeable in the test set unless the sample size is huge, because, just by chance, some site effects will be overestimated or underestimated. Also, due to the sampling variability in the test set, the differences in averages and standard deviations of pseudo-z-scores across sites cannot be exactly zero.

**Table 3a:** AIC and BIC derived measures for different models of sex.

| *model* | *AIC* | $\Delta AIC$ | *AIC.weights* | *BIC* | $\Delta BIC$ | *BIC.weights* |
| --- | --- | --- | --- | --- | --- | --- |
| M3a | -32757.25 | 0 | 0 | -32592.1 | 0 | 0 |
| M3b | -40508.33 | 0 | 0 | -39978.980 | 0 | 0 |
| M3c | -40509.24 | 0 | 0 | -39985.770 | 0 | 0 |
| M3d | -40869.51 | 0.77 | 0.77 | -39975.040 | 0 | 0 |
| M3e | -40867.07 | 0.23 | 0.23 | -40075.471 | 1 | 1 |

**Table 3b:** quality of a model fit in the test set.

| *model* | *logscore* | *skewness* | *kurtosis* | *W* |
| --- | --- | --- | --- | --- |
| M3a | 0.77 | 0.1 | 0.03 | 1 |
| M3b | 0.92 | -0.01 | -0.06 | 1 |
| M3c | 0.92 | -0.01 | -0.06 | 1 |
| M3d | 0.91 | 0.07 | 0.13 | 1 |
| M3e | 0.91 | 0.05 | 0.06 | 1 |

## Model fit comparison

In this section, we empirically compare the performance of Gaussian models and non-Gaussian models in 150 cortical thickness ROI in the UK Biobank dataset in order to quantify the potential benefit of GAMLSS modeling in a real dataset. We split the dataset into 70% training set and 30% test set. In each ROI, we fit two models, a Gaussian model

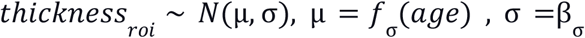

to a non-Gaussian model

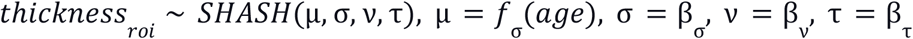

In the test set, we compare the logarithmic score and Shapiro-Wilks W, which measures the normality of pseudo-z-scores, how well are the estimated centiles calibrated. Furthermore, we also calculate skewness and kurtosis of pseudo-z-scores. Results can be seen in figure 5. According to the logarithmic score and a calibration measure W, the non-Gaussian model performed better in a vast majority of ROIs.

**Figure 5:**
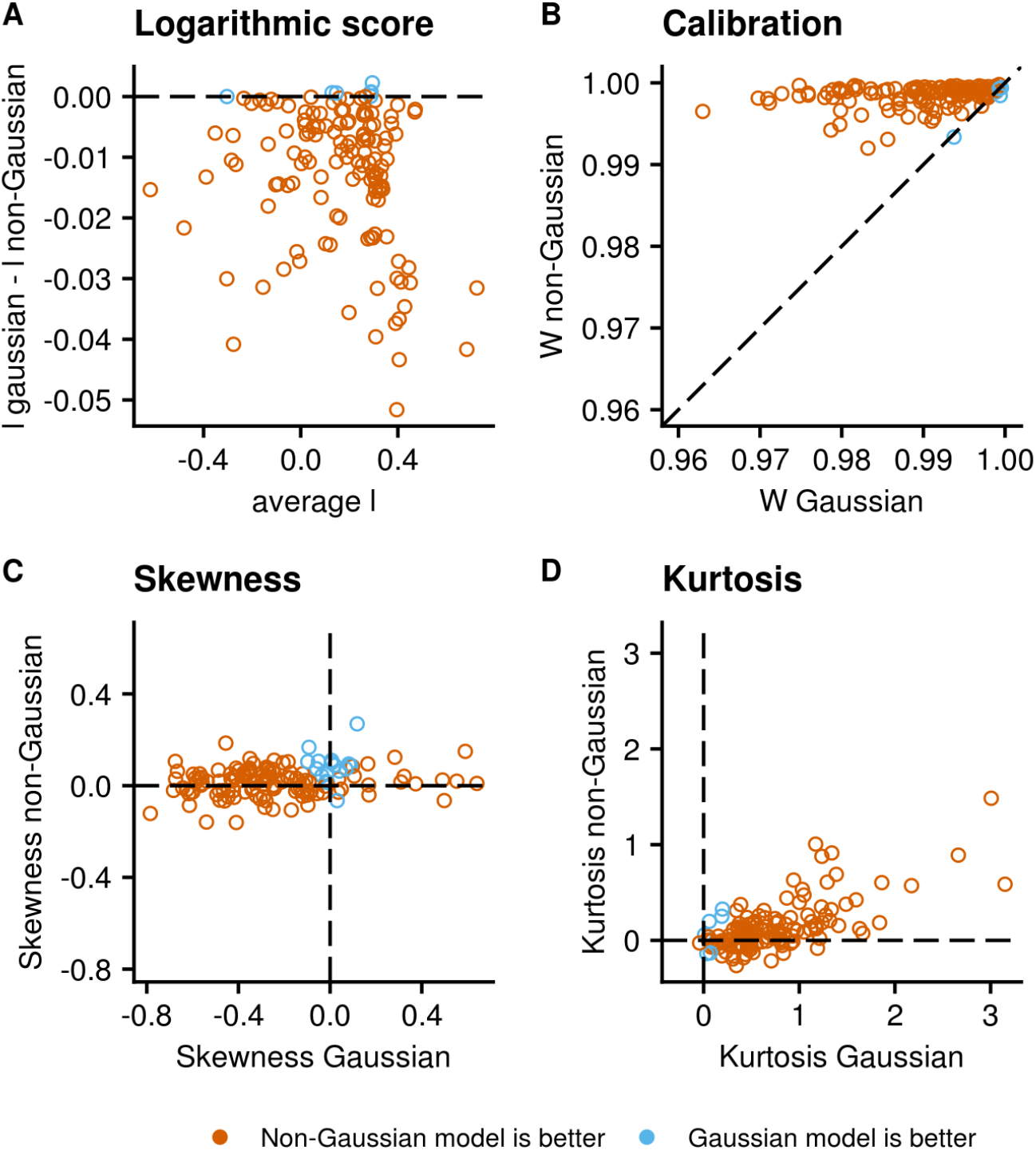
Comparison of the performance of Gaussian and non-Gaussian models in the freesurfer measures of cortical thickness in the UK Biobank dataset. Each point represents one ROI. Non-Gaussian models performed better than the gaussian model in most ROIs. **A:** Bland-Altman plot of logarithmic scores. On the x-axis is an average score of the Gaussian and non-Gaussian models. On the y-axis is their difference. **B:** Comparing models calibration measured as the Shapiro-Wilk W statistics that measure the deviation of the empirical distribution from normality. A perfect model would have the Shapiro Wilk W of 1. C, **D:** skewness and kurtosis of randomized residuals. **A:** perfect model would have skewness and kurtosis equal to zero. Deviations of skewness and kurtosis from zero and W from 1 indicate that the conditional distribution of an ROI given age was not well approximated by a given model.

## Code availability

Vode used to produce the analysis in this paper is available at https://github.com/dinga92/gamlss_normative_paper. Datasets are available from their respective repositories, some are freely available others are available with an application procedure. We provide simulated data so that all analyses could be executed and examined without any additional requirements.

## Conclusion

We introduced GAMLSS as a tool for normative modeling of brain data and described tools to diagnose and validate normative models. GAMLSS provides a powerful and flexible framework suitable to address many issues that arise while doing normative modeling. We showed several options for modeling site effects, age effects, and sex effects, including modeling of non-linear effects, heteroskedasticity, and non-gaussian distributions. Furthermore, we showed how GAMLSS models could be transferred to datasets collected at scan sites that were not available during the model fitting. GAMLSS provides a lot of modeling flexibility to researchers, therefore we described several model validation and diagnostics tools that can be used to diagnose problems with a model and decide if a model is appropriate for the data.

There is not a perfect model that would be the best for all situations. For normative modeling of brain data based on age, sex, and scan-site, the following model might be a good baseline that should perform reasonably well in most situations:

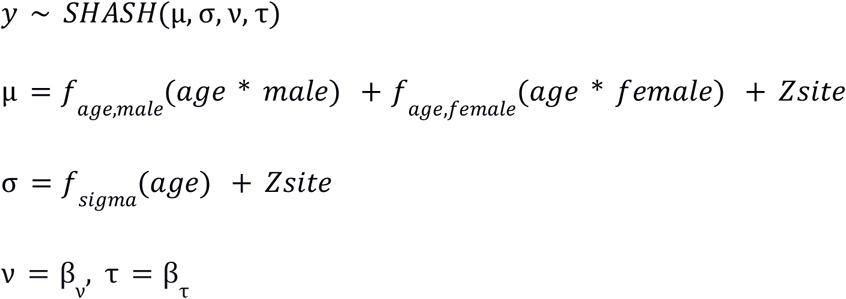

i.e., there are different age effects for men and women, the site is modeled as a random intercept, heteroskedasticity is modeled with respect to age and a site, and shape parameters are estimated from the data but do not depend on other variables. This model just a rough starting point and not a model that needs to be used in all situations. We have shown how this model can be extended to more flexibly model effects of demographic and clinical variables on brain data in the context of normative modeling.

The appropriateness of a specific modeling choice can be evaluated using the tools we presented in this paper. In general, it is reasonable to expect that age trajectories between men and women differ, as shown in many life chart studies (ref). This might not be necessary for every modality. Sex differences will be bigger for volumetric measures but smaller for cortical thickness or some functional measures. It is also reasonable to expect a non-linear effect of age on many variables, although again, this might not be necessary, especially if the normative modeling is performed in a dataset with a small age range or if a variable does not have a strong relationship with age to begin with. Biologically, we would not expect that people scanned at different scan-site will have trajectories with a different shape, however, scan-sites do have a strong effect on many data modalities, therefore, we only model site as a random intercept. However, some data modalities such as those used in tensor-based morphometry (i.e. the Jacobian determinants of a deformation field) anecdotally show only small site effects. In such cases, it might be possible to drop site effects from the model altogether. We observe that site significantly affects the variance of the data, especially for cortical thickness and volumetric measures; therefore, the model also models sigma as a random effect of site. Heteroskedasticity with respect to age is much less conclusive (Kia et al. 2020). For most of the lifespan, the variance seems to be reasonably constant, except for extremely young and extremely old people. However, the variance of some variables, such as white matter hyper-intensities, shows a strong dependence on age (Fraza et al. 2021).

The benefit of non-Gaussian modeling might be large or very small, depending on the response variable being modeled. In a large dataset such as the UK Biobank dataset with thousands of subjects in the test set, we could detect improvement of model predictions in a majority of IDPs, even in IDPs that visually look normally distributed. A similar observation was already reported by (Fraza et al. 2021). The difference in estimated z-scores will be most noticeable in the tails of a distribution, but not for relatively typical subjects. The tails contain a relatively small proportion of data, however, these are also subjects that are usually most interesting for normative modeling. For example, for median cortical thickness, which would be considered reasonably normally distributed, subjects that have a z-score 2.8 according to a Gaussian model, have a z-score 2.2 according to a non-Gaussian model (figure 4b). This difference is relatively large, and it might be even clinically important. It will also add up across multiple IDPs and affect subsequent inference, although it affects only a minority of subjects.

The limitation of this approach is that it depends on a parametric description of the distribution of the response variable, which might not always fit the data well. Especially multi-modal distributions might not be well approximated. Another limiting factor is that, in general, GAMLSS assumes that the parameters of the response distribution change smoothly as a function of input variables. If there is a sudden change of the distribution, this will not be modeled well without adjustments to the model. Last, these models are not suitable for small datasets since the higher flexibility of a model would be detrimental and might lead to overfitting.

